# Inflammation reshapes epidermal and leukocyte differentiation resulting in altered glucocorticoid responsiveness

**DOI:** 10.64898/2026.10.26.761887

**Authors:** Hildert Bronkhorst, Thomas Schrauwen, Anne Wijnants, Natalia Escobar, Susan Kloet, Annemarie H. Meijer, Marcel J.M. Schaaf, Erin Faught

## Abstract

Glucocorticoids are widely prescribed anti-inflammatory drugs that act on nearly all cell types due to the ubiquitous expression of the glucocorticoid receptor (GR). While the molecular mechanisms of GR activation have been studied extensively, it is less well understood how GR signaling modulates cell type-specific responses during inflammation. Using the unique attributes of the zebrafish, we performed single-cell RNA sequencing to characterize GR-mediated transcriptional regulation under conditions of wounding-induced inflammation. We found specific cell populations that occurred only under inflammatory conditions within epidermal cells, macrophages and neutrophils. The inflammation-specific subset of epidermal cells showed elevated expression of inflammatory markers, and these cells exhibited a reduced transactivation response to dexamethasone. This reduction in GR activation was also observed in the inflammation-specific subset of macrophages, but not in neutrophils, which retained dexamethasone responsiveness. By correlating the dexamethasone responsiveness with transcriptional coregulator expression, we identified candidate regulatory mechanisms underlying differential GR responsiveness across cell types. Taken together, we provided unique mechanistic insights into cell differentiation during a local inflammatory response and cell type-specific glucocorticoid responsiveness *in vivo*.

## INTRODUCTION

Glucocorticoids are some of the most commonly prescribed therapeutics for the treatment of a wide variety of inflammatory disorders. Approximately 3% of individuals in Western countries receive systemic glucocorticoid therapy annually, increasing to over 10% among the elderly (Laugesen et al., 2019). Despite more than seventy years of clinical use, these drugs continue to be the subject of intensive research, primarily aimed at addressing the significant adverse effects associated with long-term therapy(Koshi et al., 2022; Pofi et al., 2023). Over the past decades, substantial research efforts have been made to dissociate the beneficial anti-inflammatory effects of glucocorticoids from their deleterious side effects at the molecular level. However, these efforts have thus far met with limited success, underscoring the need for a deeper mechanistic understanding of glucocorticoid signaling (Buttgereit et al., 2025; Lesovaya et al., 2022).

Glucocorticoids exert their effects through binding and activation of the glucocorticoid receptor (GR). In the absence of a ligand, GR resides in the cytoplasm in an inactive state, stabilized by chaperones and immunophilins. Upon binding endogenous cortisol or synthetic agonists such as dexamethasone (DEX), GR undergoes a conformational change and translocates to the nucleus. There, it functions as a transcription factor, regulating the expression of thousands of genes (Lockett et al., 2024; Weikum et al., 2017). GR modulates gene expression through multiple mechanisms, including direct binding to glucocorticoid response elements (GREs) in the DNA, or indirectly through interactions with other transcription factors (Lockett et al., 2024). Efficient transcriptional regulation by GR depends on the recruitment of coactivators and corepressors, such as NCoA1, NCoA2, NCoA3, and NCoR1, as well as mediator and chromatin-remodeling complexes, such as the SWI/SNF complex (Fadel et al., 2023).

A major challenge in the study of glucocorticoids lies in the pleiotropic effects of GR signaling. GR is expressed in nearly all cell types, and its effects show large context-dependent differences between cells (Whirledge and DeFranco, 2018). The cell type-specific actions of GR can be explained by several factors, including the levels of cell-specific transcription factors and the availability of coregulators, which have primarily been studied in cultured cells (Love et al., 2017; Mao et al., 2023; Clarisse et al., 2024; Fadel et al., 2023). Some studies have linked GR-mediated phenotypes *in vivo* to interactions between GR and other transcriptional regulators; for example, GR activity in mouse liver or muscle is critically dependent on the levels of the transcription factor FOXO1 or the histone demethylase LSD1, respectively (Cai et al., 2024; Makris et al., 2025). However, unbiased approaches capable of assessing cell type-specific GR activity across multiple tissues simultaneously remain needed.

In the present study, we performed single-cell RNA sequencing (scRNA-seq) in zebrafish larvae to resolve cell type-specific GR activity *in vivo*. Using caudal fin wounding as a model of acute inflammation in combination with glucocorticoid treatment, we characterized GR activity and coregulator interactions under inflammatory conditions at single-cell resolution. Zebrafish provide an excellent system for studying GR action and inflammation (Dinarello et al., 2020; Schaaf et al., 2009). The glucocorticoid signaling pathway is highly conserved between humans and zebrafish, and their GRs show strong structural and functional similarity(Alsop and Vijayan, 2008; Schaaf et al., 2008). Inflammation can be induced via caudal fin wounding, which is typically performed in early larval stages (2-5 days post-fertilization (dpf)). This injury elicits a local inflammatory response characterized by recruitment of macrophages and neutrophils as well as inflammatory gene expression, both of which are attenuated by glucocorticoid treatment (Mathew et al., 2007; Chatzopoulou et al., 2016; Xie et al., 2021). Previous zebrafish scRNA-seq studies following caudal fin wounding have mapped regeneration processes in considerable detail, focusing on time points from 1 day post-amputation onwards (Chen et al., 2025; Hou et al., 2020; Laplace-Builhé et al., 2021; Sinclair et al., 2021; Tang et al., 2022). However, early time points that are critical for capturing acute inflammatory activity, as well as the effects of glucocorticoid treatment on injured tissue, are still to be explored.

Therefore, in the present study, we examined the transcriptomic effects of vehicle (DMSO) and DEX treatment by scRNA-seq and complementary bulk RNA-seq at 4 hours post-amputation (hpa), a time point coinciding with peak inflammatory activity (Chatzopoulou et al., 2016; Xie et al., 2019). We identified specific populations of epidermal and leukocyte cells that are reprogrammed upon caudal fin wounding, and further characterized these key cell types in terms of GR activity. Importantly, these populations exhibited distinct glucocorticoid responsiveness and a specific profile of GR transcriptional coregulator expression. Together, these datasets provide a comprehensive overview of cell type-specific glucocorticoid responsiveness during acute inflammation and early regeneration *in vivo*.

## RESULTS

### Inflammatory and metabolic pathways are affected by zebrafish caudal fin amputation

Using established transgenic zebrafish lines (*Tg (mpeg:mCherry/mpx:eGFP)*), in which macrophages and neutrophils are fluorescently labelled, we first assessed leukocyte localization at a wound site and whether this could be modulated by DEX. To this end, 3 dpf larvae underwent caudal fin amputation just posterior to the notochord after 2 hours of pretreatment with DMSO or dexamethasone (DEX) (Fig.1a,b). Treatment with DEX reduced the number of neutrophils that were localized to the wound by half compared to vehicle treated larvae (p<0.0001; Fig 1c). Similarly, in macrophages, DEX also reduced the number of neutrophils that migrated towards the wound by 20% (p<0.001; Fig 1d). These results are consistent with other reports in zebrafish in which other glucocorticoids can also reduce the number of migrating leukocytes (Faught and Schaaf, 2023; Xie et al., 2019).

**Figure 1.**
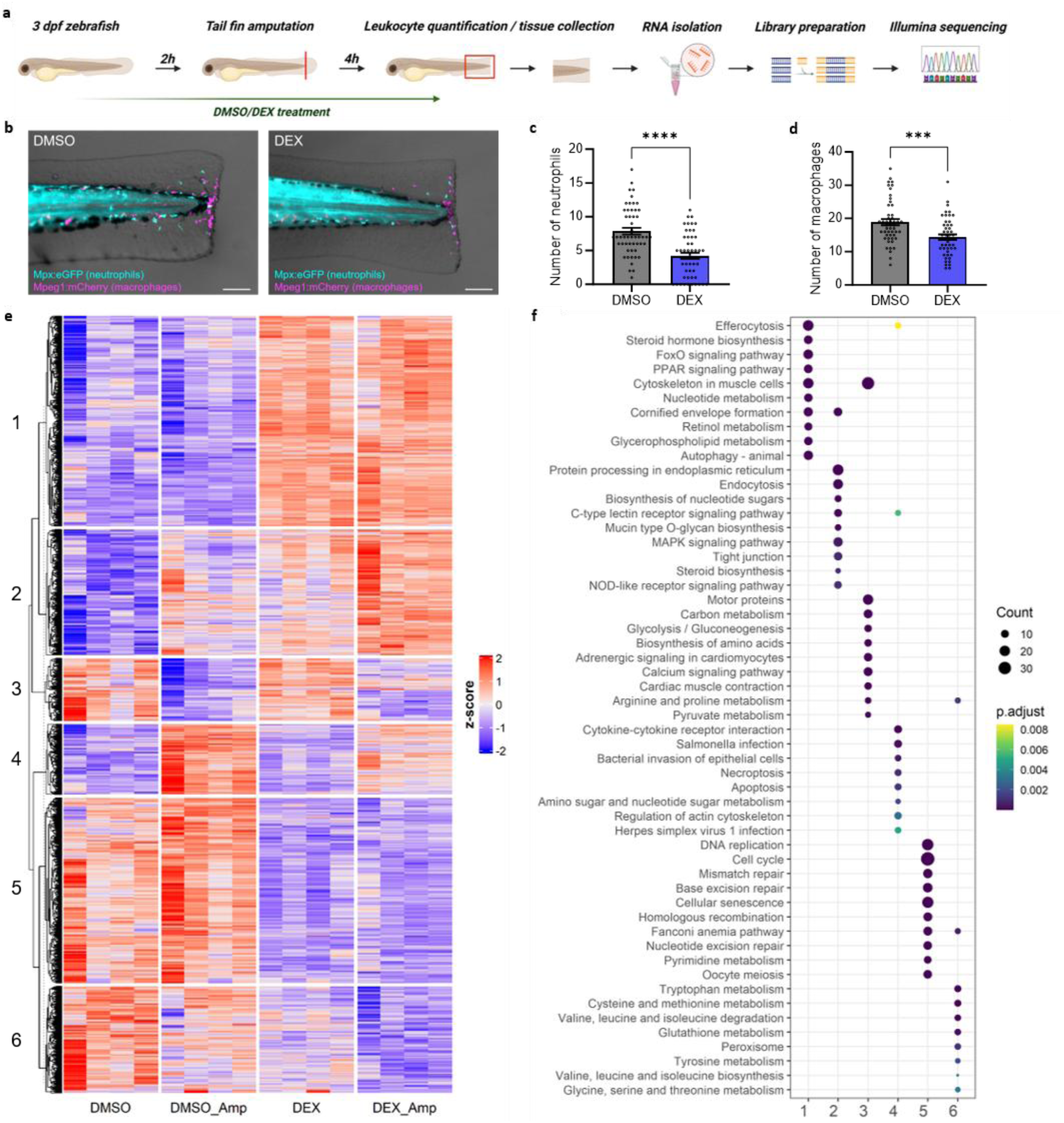
Inflammatory and metabolic pathways are affected by wounding. (a) Schematic of the experimental setup for bulk RNA-seq on zebrafish caudal fin tissue. (b) Representative images of macrophage and neutrophil localization at the wound site *Tg (mpeg1.1:mCherry/mpx:eGFP)* larvae. (c) Quantification of the number of neutrophils (GFP-positive cells) within 200µm of the wound site. (d) Quantification of the number of macrophages (mCherry-positive cells) within 200µm of the wound site. Statistical significance in (c) and (d) was determined with a Student’s t-test (p<0.05; *** p<0.001, ****p<0.0001) (e) Heatmap of all genes from the four biological replicates differentially expressed upon amputation or DEX treatment with adjusted p-value <0.05 in at least one contrast. (f) Top 10 pathways ranked by significance for each gene cluster identified in Fig.1e.

To obtain statistically robust, in-depth insight into transcriptional responses to wounding and glucocorticoid treatment, we next performed bulk RNA-seq on zebrafish caudal fin tissue. At 4 hpa, caudal fin tissue was collected, and RNA was extracted for sequencing (Fig.1a). We assessed gene expression across four experimental groups: DMSO-treated non-amputated (DMSO), DMSO-treated amputated (DMSO_Amp), DEX-treated non-amputated (DEX), and DEX-treated amputated (DEX_Amp). Differentially expressed genes (DEGs) were identified across multiple pairwise comparisons. Amputation (DMSO_Amp versus DMSO) resulted in 583 significant DEGs, while DEX treatment (DEX versus DMSO) gave 1291 DEGs. The combined effect of amputation and DEX treatment (DEX_Amp vs DMSO) yielded 2095 DEGs. In addition, 1588 DEGs were detected between DEX_Amp and DMSO_Amp, reflecting DEX-specific effects in the amputated condition (Fig.1e, Table S1). Thus, DEX treatment resulted in a higher number of DEGs than amputation, and the combined DEX_Amp treatment exhibited the largest transcriptional response, indicating an additive effect of amputation and DEX treatment.

All significant DEGs were used to generate a heatmap to investigate global regulatory patterns (Fig.1e, Table S2). Unsupervised clustering identified six major gene clusters with distinct regulation across the four conditions. To gain insight into the pathways affected by amputation and DEX treatment, we performed pathway analysis for all gene clusters identified in the heatmap. The top 10 significant pathways for each cluster are shown in Fig.1f and the full analysis is presented in Table S3. Cluster 1 and 5 contained genes that were respectively induced or repressed by DEX treatment independent of amputation. Cluster 1 comprised pathways related to metabolic rewiring, steroid processing, and anti-inflammatory signaling, and includes multiple canonical GR target genes such as *fkbp5*, *pck1*, and the anti-inflammatory genes *tsc22d3* and *dusp1*. In contrast, cluster 5 showed enrichment of cell proliferation and DNA replication pathways, involving genes like *mcm8*, *pola1*, *ccna2*, and *msh2*. Cluster 2 and 6 showed genes that were respectively induced or repressed by both amputation and DEX treatment, resulting in the most pronounced expression changes in the DEX_Amp condition. Cluster 2 included processes for protein processing and secretion, barrier formation, and pattern recognition receptor signaling, with representative genes such as *nfkbiaa*, *il1b*, *lamc2*, and *hsp90aa1*. These pathways are likely activated by amputation and DEX to control inflammation and maintain tissue integrity. Analysis of cluster 6, with genes such as *aox5*, *bcat2*, *dnmt1*, and *oplah*, revealed a set of redox and antioxidant defense pathways, indicating amputation and DEX-induced suppression of oxidative stress-related metabolic programs. Finally, genes in cluster 3 and 4 were respectively strongly repressed or induced by amputation, with both responses attenuated by DEX. Cluster 3 contained multiple metabolic processes, as well as calcium and IgSF CAM signaling, with contributing genes *pkmb*, *ckma*, *dcc*, and *p2rx8*. The regulatory pattern across the treatments suggests that homeostatic metabolism, as well as neuromuscular development, are suppressed following injury, possibly to prioritize host defense and tissue repair. This suppression is attenuated by DEX treatment. Cluster 4 is enriched for classical innate immune activation pathways, including cytokine signaling, host-pathogen interaction, cell migration, cell death and clearance, and tissue repair signaling. Among the most strongly regulated genes in this cluster were *cxcl18a.1*, *il11*, *lepb*, *gcga*, *ednrb*, *mmp9*, *cd44a*, and *lsp1*. This cluster reflects the classical anti-inflammatory effects of DEX, but also includes DEX-mediated suppression of other strongly amputation-induced processes such as tissue repair.

Collectively, the bulk RNA-seq results show that DEX modulates almost all transcriptional responses to amputation, attenuating strong regulation upon amputation (clusters 3 and 4) and enhancing weaker responses (clusters 2 and 6). In contrast, strong regulation by DEX remains largely unaffected by amputation (clusters 1 and 5). Interestingly, no substantial gene cluster was identified that was solely regulated by amputation and unaffected by DEX.

### scRNA-seq analysis reveals an amputation-specific epidermal cell population

Because bulk RNA-seq can mask cellular heterogeneity of gene expression, we applied single-cell RNA-seq (scRNA-seq) on zebrafish caudal fin tissue under the same treatment conditions (Fig.2a-c). After quality control and preprocessing, 76,815 high-quality cells were retained for downstream analysis. Clustering (Fig.2b) followed by manual annotation based on marker genes (Fig.2d, Table S4) identified all major cell types present in the zebrafish caudal fin, including innate immune populations such as neutrophils and macrophages. Next, integration of the data from the four experimental groups into a single dataset enabled direct comparison of treatment effects across conditions. The integrated dataset faithfully recapitulated upregulation of pro-inflammatory genes following amputation and its transrepression by DEX (e.g., *cxcl18a.1*, Fig.2c), as well as activation of canonical glucocorticoid-induced genes (e.g., *fkbp5*, Fig.2c). Accordingly, we used this dataset to investigate amputation- and DEX-dependent transcriptional changes at single-cell resolution.

**Figure 2.**
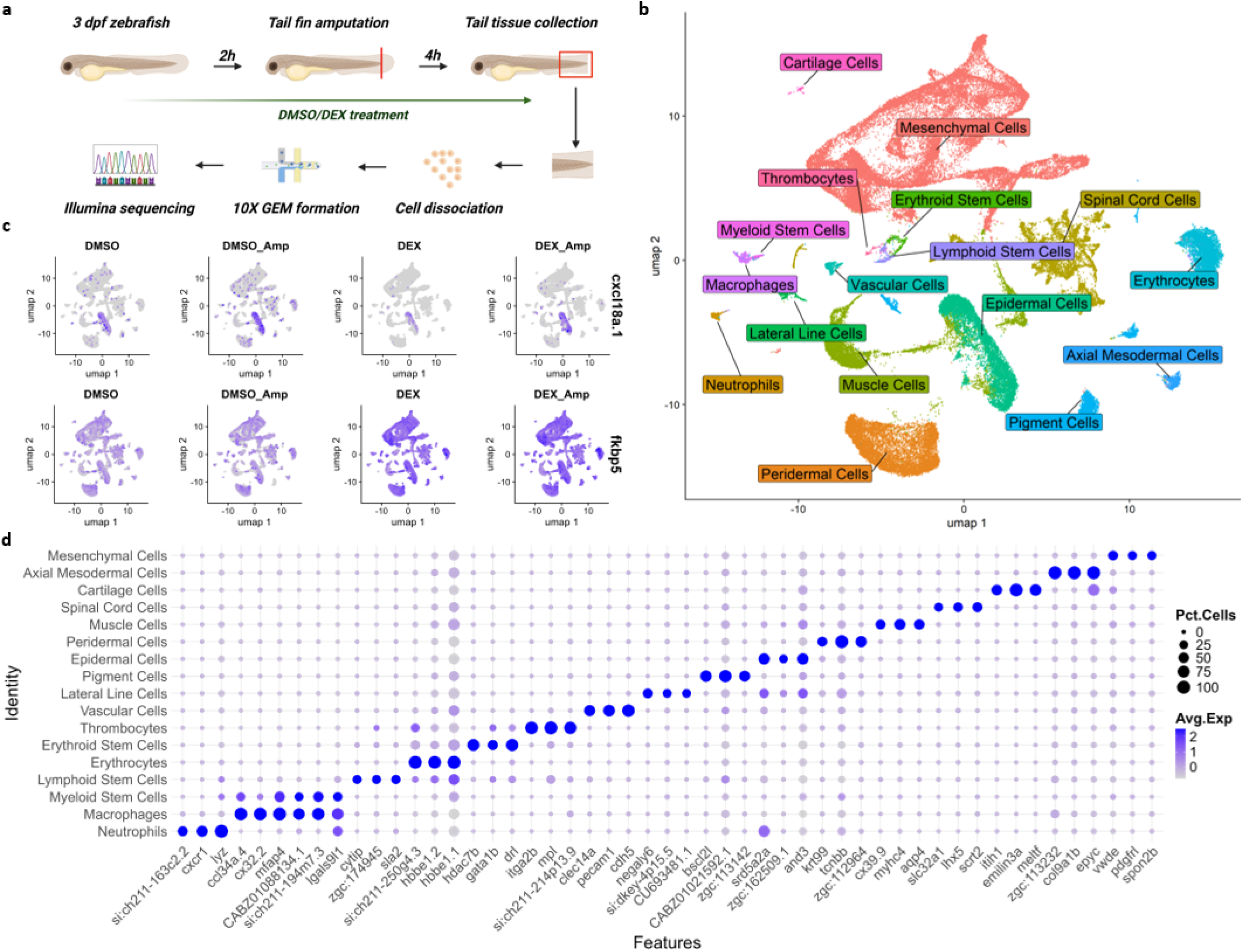
Overview of the scRNA-seq experiment. (a) Schematic of the experimental setup for scRNA-seq on zebrafish caudal fin tissue. (b) UMAP projection of identified tissues and cell types of interest. (c) Feature plot of *cxcl18a.1* and *fkbp5* showing successful RPCA integration of the four experimental conditions and the expected impact of wounding and DEX treatment. (d) Dot plot of the top 3 marker genes for the tissues and cell types shown in (b).

The preceding bulk RNA-seq analysis identified genes that were strongly upregulated following amputation (Fig.1b). Mapping their expression in the scRNA-seq dataset revealed that these genes were highly enriched in the epidermal cell cluster (Fig.S1). Consequently, we isolated the epidermal cells from the scRNA-seq data and reclustered them to resolve subpopulations (Fig.3a-d, Table S5). Strikingly, a small epidermal subcluster emerged in the two amputation conditions (DMSO_Amp and DEX_Amp), but was absent from both non-amputated groups (Fig.3a). This cluster comprised 53 of the 1597 epidermal cells (3.3%) in DMSO_Amp, and 53 of the 2388 epidermal cells (2.2%) in the DEX_Amp condition. This amputaion-specific subcluster belonged to a broader caudal fin population characterized by expression of *mxra8b*, *krtSc*, and *krtS7*, but was distinguished from other epidermal cells by a set of highly specific marker genes such as *fam167b* and *fstl3* (Fig.3b,d). These markers include several of the most strongly upregulated genes identified in the bulk RNA-seq analysis (Fig.1b,c), such as *gcga*, *adam8a*, *fstl3*, *cd44a*, and *lepb*. In addition, these amputation-specific markers include genes that are implicated in inflammation and tissue remodeling (e.g., *tnfb*, *mmp* family members, *il1b*, and *lect2l*), as well as genes encoding metabolic hormones (e.g., *gcga* and *fstl3*) (Fig.3b, Fig.S2). To functionally characterize the amputation-specific cell population, we performed Gene Ontology (GO) analysis on the amputation-specific marker genes (Fig.3c, Table S6). The most significant GO terms were associated with the response to wounding, regeneration, growth, development, and signal transduction. Finally, we inferred transcription factor activity in amputation-specific cells and found high activity from inflammatory and proliferative transcription factors encoded by genes such as *sp2*, *fos1a*, *ets2*, and *rela* that show 2-14 fold higher activity in amputation-specific cells than in the other epidermal cells (Fig.3e). Together, these findings show that the major transcriptional responses following caudal fin amputation are concentrated to a small subset of epidermal cells. While DEX treatment did not abolish the formation of these amputation-specific cells, it reduced their relative abundance by 30%. Amputation-specific cells display a pronounced inflammation- and regeneration-associated transcriptional profile. We therefore propose that this population plays a key role in mediating inflammatory signaling at the wound site, and in initiating tissue remodeling processes required for fin regeneration.

**Figure 3.**
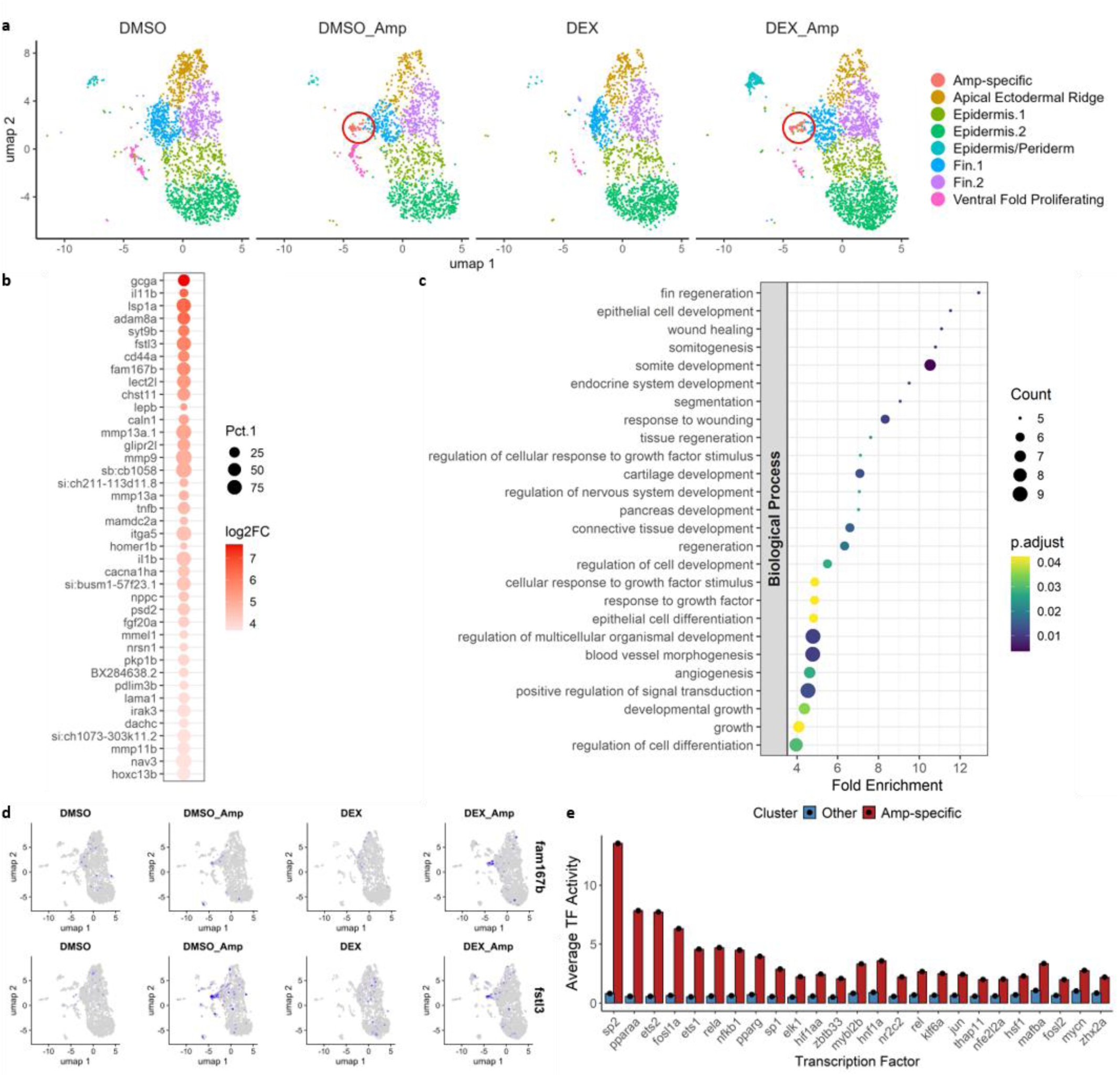
A distinct epidermal cell population arises after wounding. (a) UMAP projection of the epidermal cell subset reveals an amputation-specific cluster in both amputation conditions. (b) Top 40 marker genes by log2 fold change of the Amp-specific cluster versus all other epidermal cells. (c) Gene ontology of the Amp-specific marker genes. (d) Feature plots with the 2 most significant marker genes of the Amp-specific cluster. (e) Transcription factor activity in Amp-specific epidermal cells and other epidermal cells. Shown are 25 transcription factors with the largest difference between the two cell types.

### Amputation-specific epidermal cells are less responsive to DEX-induced gene upregulation

Having identified the epidermal cell population most strongly affected by inflammation, we next examined its responsivity to DEX. We defined a GR Module Score (GMS) as the normalized mean expression of the top 100 epidermal DEX-responsive genes per cell, providing a proxy for the level of GR transcriptional activity in each cell. We then compared mean GMS values between amputation-specific epidermal cells and all other epidermal cells. The GMS for upregulated genes was significantly lower in amputation-specific cells than in other epidermal cells (Fig.4a), suggesting that GR transactivation was curtailed in amputation-exposed epidermal cells. In contrast, no difference in GMS was observed for downregulated genes in the DEX_Amp condition (Fig.4b), meaning that repression was equally effective across both cell types. We subsequently determined which of the DEX-upregulated genes contributed to the reduced GMS in amputation-specific cells (Fig.4c, Table S7). Genes showing reduced induction included multiple stress-related transcriptional regulators (*klf2a, klf9*, *tob1b*, *igfbp5b*, and *dusp1*), as well as genes involved in epithelial adhesion (*itga6b*, *itgb4*, *thy1*, *pof1b*, and *mtss1la*). These findings indicate that specific and important gene programs in amputation-specific cells are less responsive to DEX in these cells, resulting in an overall reduction in the GMS.

**Figure 4.**
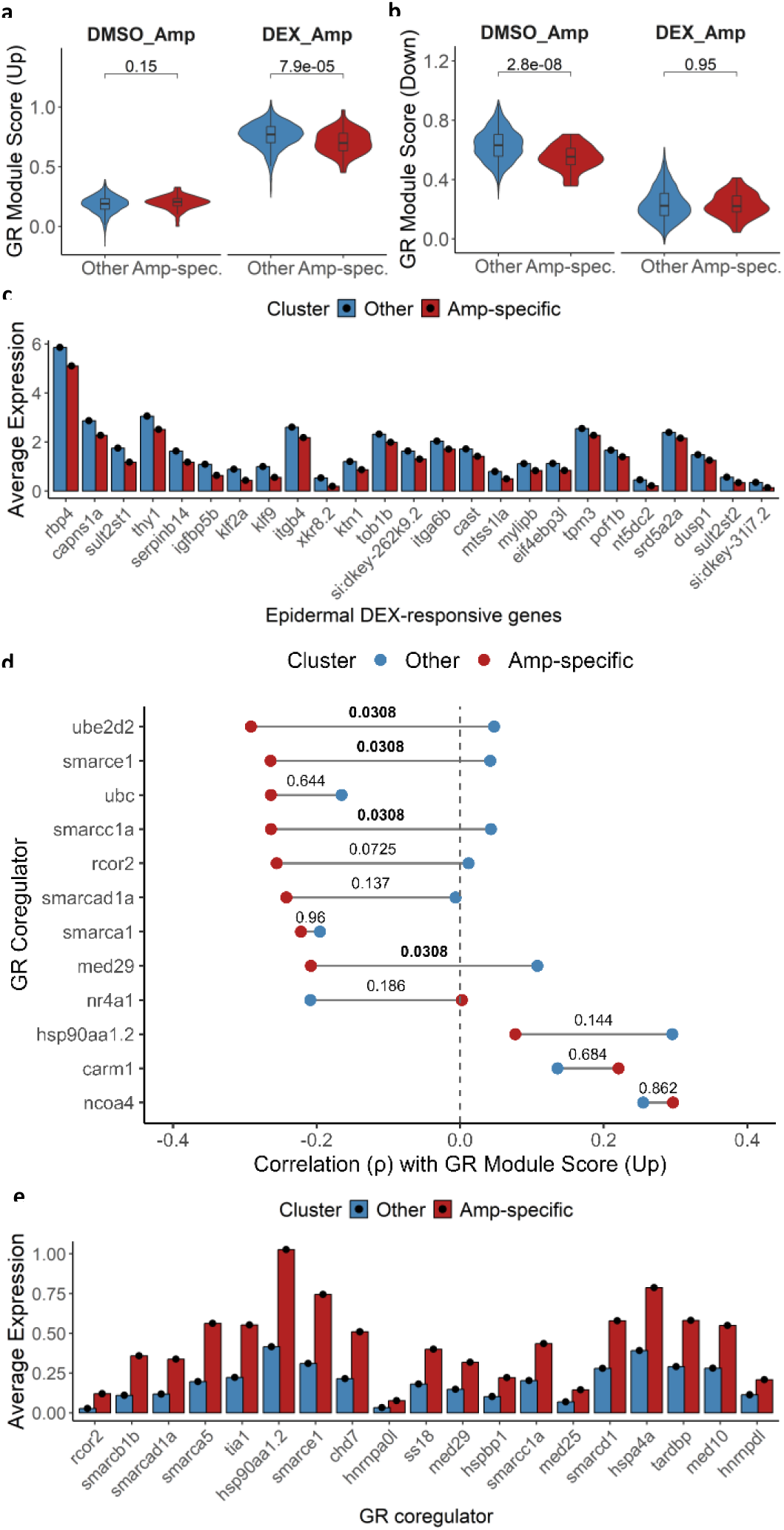
Amputation-specific epidermal cells are less responsive to DEX. (a-b) GR Module Score with up-(a) or downregulated (b) genes in the Amp-specific and the other epidermal cell clusters. Statistical significance between the two clusters in each condition was assessed using the Wilcoxon test as indicated. (c) Top 25 DEX-responsive upregulated genes with the largest reduced expression differences between the Amp-specific and other epidermal cell clusters. (d) Spearman correlations between GR coregulator expression and the upregulated GR Module Score in Amp-specific and other epidermal cell clusters, restricted to correlations with |ρ| > 0.2. The numbers indicate the adjusted p-value of the statistical difference between the two correlation values determined by a Fisher Z-transformation and Benjamini-Hochberg procedure. (e) Differential GR coregulator expression in Amp-specific and other epidermal cells with |log2 fold change| > 2.

To investigate the mechanism underlying the reduced responsiveness to GR-induced gene activation in the amputation-specific cells, we correlated the GMS with the expression of GR transcriptional coregulators at single-cell resolution for both amputation-specific and other epidermal cells (Fig.4d). Significant differences in correlation between the two cell types were observed for *ube2d2*, *smarce1*, *smarcc1a*, and *med2S*. The correlation between the expression of these coregulators and the GMS was strongly negative in amputation-specific cells, which means that the reduced DEX responsiveness in these cells is associated with increased expression of these coregulators, suggesting a negative impact of these factors on GR-induced gene transcription. Consistent with this result, differential expression analysis of GR coregulators revealed that many coregulators with a negative correlation were significantly upregulated in amputation-specific cells compared with other epidermal cells (Fig.4e). These include components of the SWI/SNF complex and related chromatin remodelers (*smarcb1b*, *smarce1*, *ss18*, *smarcc1a*, *smarcd1*, *smarcad1a, chd7*), a transcriptional corepressor (*rcor2*), subunits of the Mediator complex (*med29*, *med25, med10*), and chaperones (*hsp90aa1.2*, *hspbp1*, *hspa4a*). Although the activity of these factors is context-dependent, their overexpression in amputation-specific cells, coupled with negative correlation to DEX-mediated gene induction, suggests a repressive chromatin and GR coregulator environment that limits GR-driven gene activation in amputation-specific epidermal cells.

### Identification of amputation-specific neutrophils and macrophages

Leukocytes respond to caudal fin amputation by migrating to the wound site, where they contribute to pathogen defense and tissue repair(Xie et al., 2021). To characterize these cells in more detail, we isolated the leukocyte population from our scRNA-seq dataset, which included cells identified as macrophages, neutrophils, T helper cells, and multiple stem cell types (Fig.5a). Subclustering and annotation revealed distinct macrophage and neutrophil populations that were specific to the amputation condition irrespective of DEX treatment, which were therefore designated as Amp-macrophages and Amp-neutrophils (Fig.5a, Table S8). For neutrophils, 26 out of 89 (29%) in DMSO_Amp and 31 out of 90 (34%) in DEX_Amp clustered as Amp-neutrophils. For macrophages, 47 out of 97 (48%) in DMSO_Amp and 64 out of 138 (46%) in DEX_Amp were classified as Amp-macrophages, although these were less clearly separated from other macrophage populations. Thus, the proportions of amputation-specific populations did not differ significantly between DMSO and DEX treatment.

**Figure 5.**
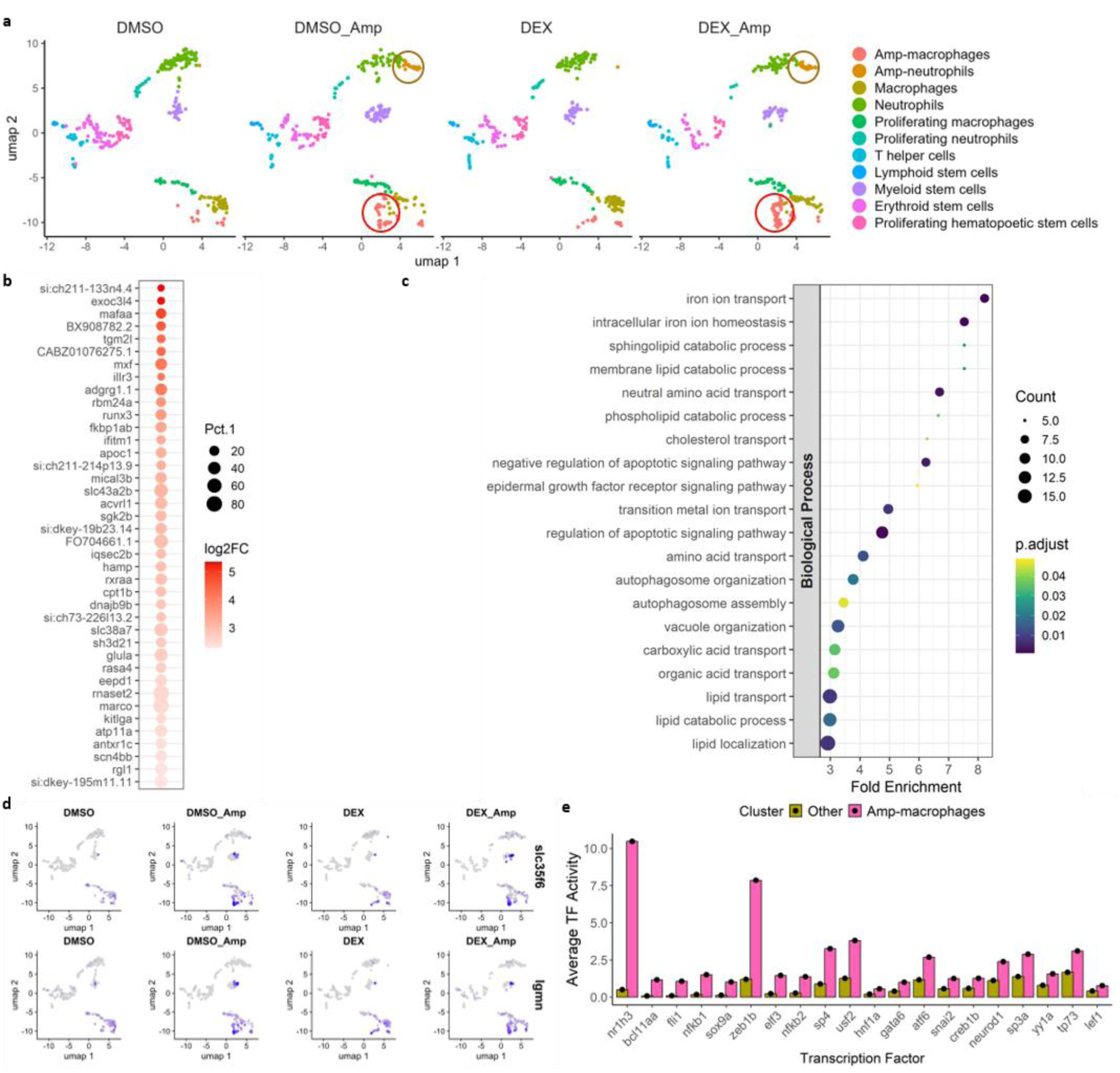
Distinct macrophage cell populations arise after wounding (a) UMAP projection of the leukocyte subset reveals amputation-specific neutrophils and macrophages in both amputation conditions. (b) Top 40 marker genes by log2 fold change for the Amp-macrophages versus other macrophages. (c) Gene ontology of the Amp-macrophage marker genes. (d) Feature plots with the 2 most significant marker genes of the Amp-macrophages. (e) Transcription factor activity in Amp-macrophages and other macrophages. Shown are 20 transcription factors with the largest difference between the two cell types.

We subsequently analyzed the marker genes and their associated gene ontology of Amp-macrophages following the same approach as used for amputation-specific epidermal cells (Fig.5b-e). Amp-macrophage marker genes and GO terms were enriched for lipid trafficking and degradation, autophagy, iron homeostasis, and regulation of apoptotic signaling (Fig.5b-c, Fig. S3a, Table S9), with the most significant marker genes *slc35fc* and *lgmn* (Fig.5d). These processes are characteristic of stress-responsive macrophages with high degradative activity of lipid material. Moreover, transcription factor activity analysis revealed a tenfold higher activity of *nr1h3* in Amp-macrophages, with *nr1h3* encoding the nuclear receptor LXRα, which drives cholesterol homeostasis and inflammatory reprogramming (Fig.5e). Amp-neutrophils exhibited a marker gene expression profile characteristic of active neutrophils, with GO terms including superoxide metabolic process, defense response to bacterium, cell chemotaxis, and inflammatory response (Fig.6a,b, Fig.S4a, Table S10). The most significant marker genes were *aif1l* and *csf2rb* (Fig.6c). In addition, Amp-neutrophils showed high activity of transcription factors regulating the immune response, i.e., *relb*, *hsf1*, *rel*, and *runx3* (Fig.6d). Overall, the marker genes and their associated GO terms of these Amp-macrophages and Amp-neutrophils are clearly indicative of activated leukocytes that have responded to tissue wounding.

**Figure 6.**
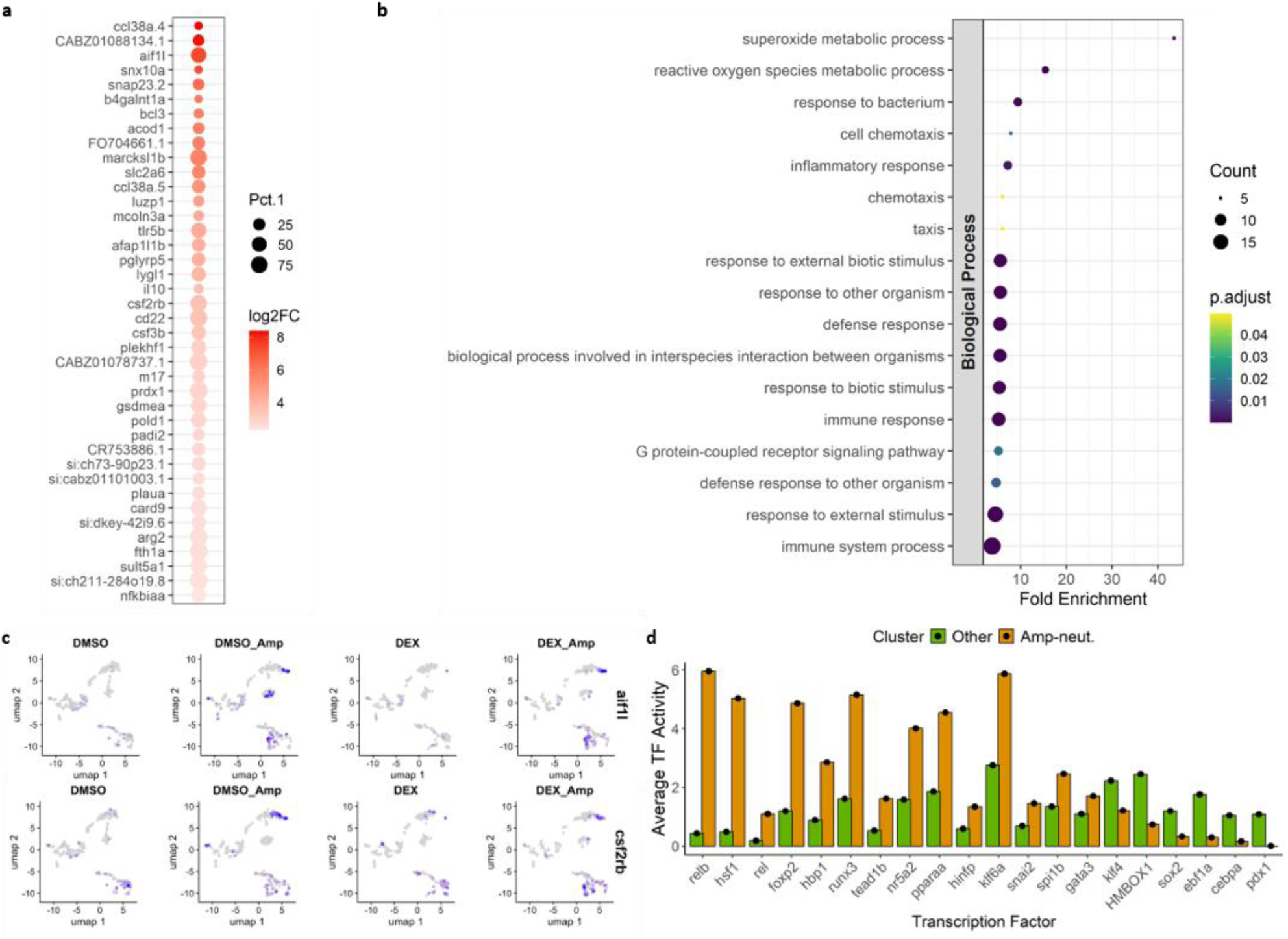
Distinct neutrophil cell populations arise after wounding. (a) Top 40 marker genes by log2 fold change for the Amp-neutrophils versus other neutrophils. (b) Gene ontology of the Amp-neutrophil marker genes. (c) Feature plots with the 2 most significant marker genes of the Amp-neutrophils. (d) Transcription factor activity in Amp-neutrophils and other neutrophils. Shown are 20 transcription factors with the largest difference between the two cell types.

### Amputation-specific macrophages are less responsive to DEX-induced gene upregulation

After the identification of Amp-macrophages and Amp-neutrophil clusters, we next investigated their responsiveness to DEX using the same approach as for the epidermal cells. First, the GMS (defined as the mean normalized expression of the top 100 DEX-responsive genes) was determined in macrophages. We found that Amp-macrophages exhibited a reduced GMS for upregulated genes compared to other macrophages (Fig.7a), while for downregulated genes the GMS did not differ significantly (Fig.7b). The genes showing reduced induction by DEX in Amp-macrophages include transcriptional regulators (*raraa*, *stk17al*, *fam107b*, *dusp2*, *trib3*, *nfil3-c, tsc22d3*), as well as proliferation-related genes (*pik3r1*, *pim1*, *rps6kb1b*, *mid1ip1l*), and genes involved in membrane transport (*slc7a3a*, *mctp2b*, *slco2a1*; Fig.7c, Table S11). The reduced expression of immunomodulatory genes such as *raraa, dusp2,* and *tsc22d3* in Amp-macrophages suggests an altered capacity to mount an anti-inflammatory transcriptional program in response to DEX treatment.

**Figure 7.**
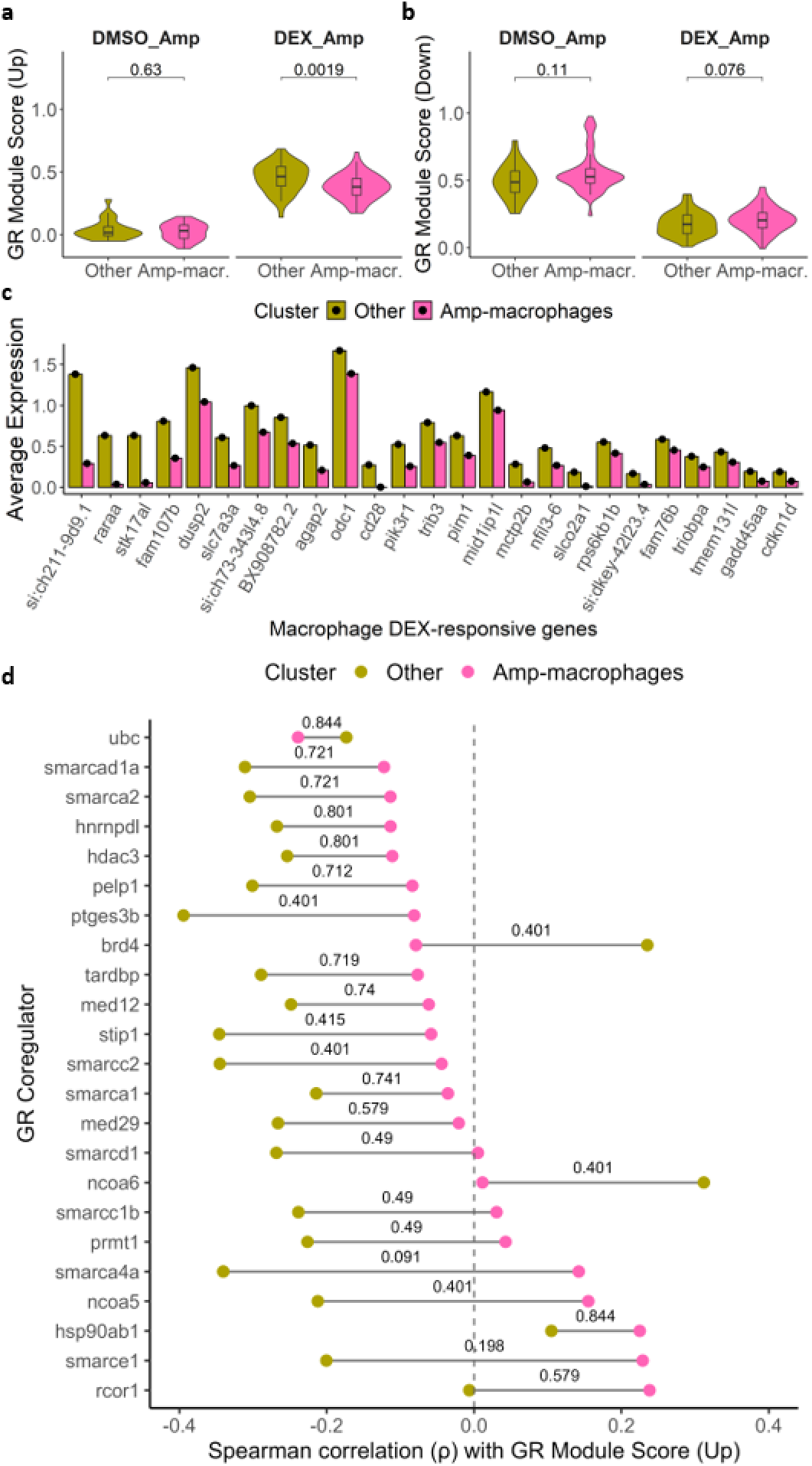
Amputation-specific macrophages are less responsive to DEX activation. (a-b) GR Module Score with up-(a) or downregulated (b) genes in the Amp-macrophages and the other macrophages. Statistical significance between the two clusters in each condition was assessed using the Wilcoxon test as indicated. (c) Top 25 DEX-responsive upregulated genes with the largest expression differences between the Amp-macrophages and other macrophages. (d) Spearman correlations between GR coregulator expression and the GR Module Score (down) in Amp-macrophages and other macrophages, restricted to correlations with |ρ| > 0.2.

Next, we correlated the GMS with the expression of transcriptional coregulators in Amp-macrophages and other macrophages (Fig.7d). In Amp-macrophages, few correlations exceeded |ρ| > 0.2, with the strongest positive correlation in Amp-macrophages observed for *rcor1*, an important transcriptional corepressor, which may contribute to the reduced GMS in these cells. The observed correlations in other macrophages were predominantly negative, and no statistically significant differences were found between correlations in Amp-macrophages and other macrophages, likely limited power due to the relatively small number of cells. Probably for the same reason, no significant differences in expression of transcriptional coregulators were observed (Fig.S3b). In short, analysis of macrophages in our dataset revealed that Amp-macrophages, characterized by a strong lipid metabolism signature, show increased expression of the corepressor gene *rcor1* and exhibit a reduced responsiveness to glucocorticoid-induced gene activation.

### Amputation-specific neutrophils retain glucocorticoid responsiveness

For the neutrophils, we observed no difference between Amp-neutrophils and other neutrophils in the GMS for upregulated genes (Fig.8a). Notably, amputation-specific neutrophils, regardless of treatment, exhibited a lower GMS for downregulated genes (Fig.8b). This suggests that amputation alters GR activity under both baseline and upon DEX-treated conditions. We subsequently plotted the downregulated genes that are responsible for the reduced GMS in Amp-neutrophils (Fig.8c, Table S12). Several genes with lower expression in Amp-neutrophils were involved in DNA transcription and chromatin organization (*nucks1a*, *histh1l*, *polr2eb*, *cebpz, psip1b*, *h1-10*), whereas others were associated with inflammation (*lgals9l3*, *ctsk*, *egr2a*, *ilf3a*), and mitochondrial processes (*aldh6a1*, *gpx1a*, *acadl*, *gstk1*, *mt-atp8).* Although many of these genes were already expressed at lower levels in amputation-specific neutrophils under the DMSO condition, DEX further reduced the relative expression level in these cells (Table S12). This finding suggests that Amp-neutrophils are still fully responsive to DEX in their inflammatory state, in contrast to the amputation-specific epidermal cells and Amp-macrophages.

**Figure 8.**
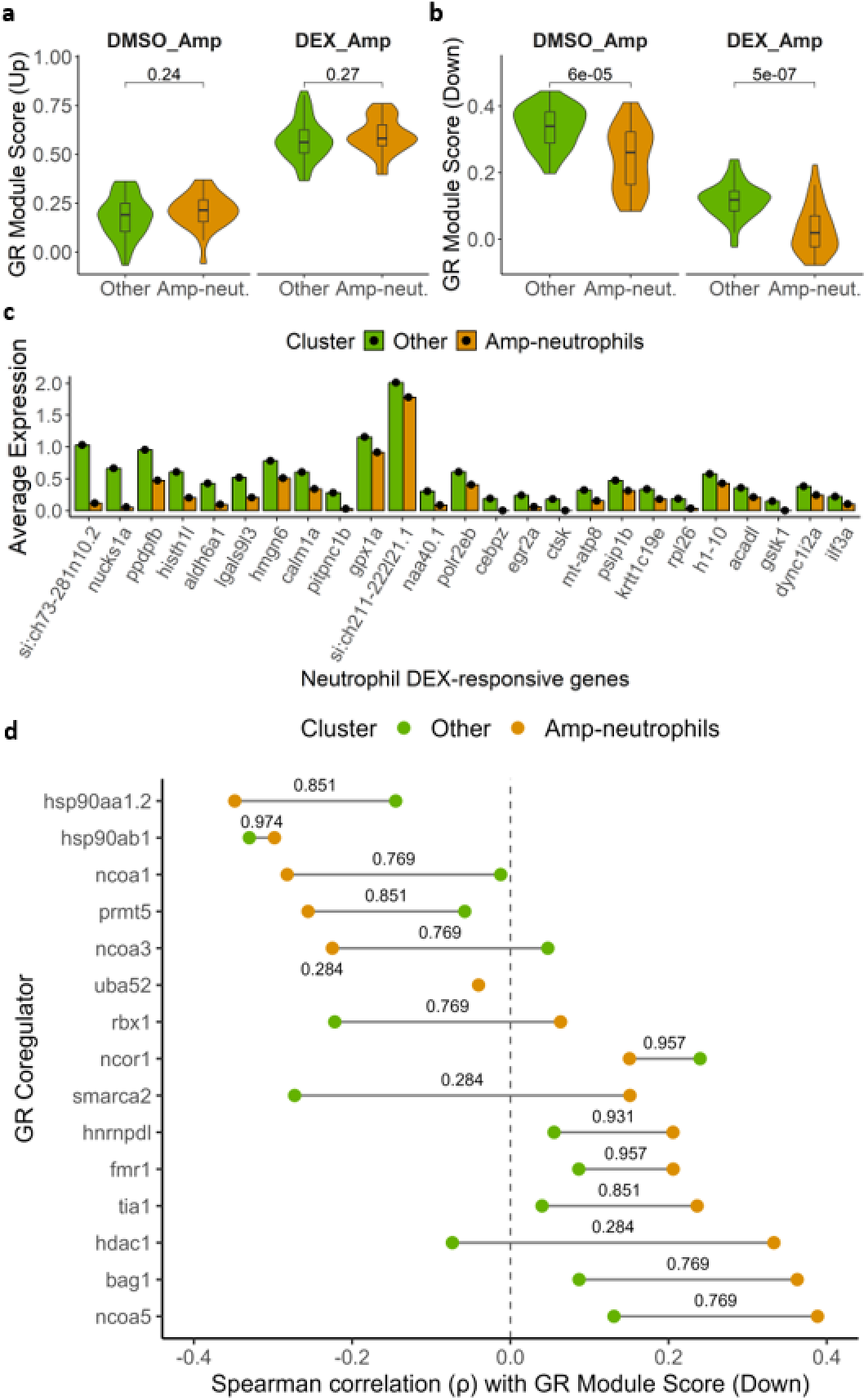
Amputation-specific neutrophils retain DEX responsivity. (a-b) GR Module Score with up-(a) or downregulated (b) genes in the Amp-neutrophils and the other neutrophils. Statistical significance between the two clusters in each condition was assessed using the Wilcoxon test as indicated. (c) Top 25 DEX-responsive downregulated genes with the largest expression differences between the Amp-neutrophils and other neutrophils. (d) Spearman correlations between GR coregulator expression and the GR Module Score (down) in Amp-neutrophils and other neutrophils, restricted to correlations with |ρ| > 0.2.

Finally, we correlated the GMS for downregulated genes with the expression of transcriptional coregulators (Fig.8d). Interestingly, a negative correlation was observed for *ncoa1* and *ncoa3* in Amp-neutrophils, whereas *ncoa5* showed a positive correlation. This suggests that lower expression of GR-repressed genes (the most differentially regulated shown in Fig.8b) is associated with higher expression of *ncoa1* and *ncoa3*, and lower expression of *ncoa5*. Similar to the macrophage analysis, no significant differences in expression of transcriptional coregulators were observed due to the low cell number (Fig.S4b). Overall, these data indicate that neutrophils retain glucocorticoid responsiveness for both up- and downregulated genes under inflammatory conditions. GR activity in Amp-neutrophils was associated with a distinct cofactor correlation profile, from which cell type-specific GR modulation can be inferred.

Taken together, our scRNA-seq analysis of epidermal cells, macrophages, and neutrophils revealed cell type-specific glucocorticoid responsiveness under inflammatory conditions. In amputation-specific epidermal cells and in Amp-macrophages, we observed a reduced transactivation response to dexamethasone, while in in Amp-neutrophils dexamethasone responsiveness was retained (Fig.9).

## DISCUSSION

In the present study, we exploited a well-characterized larval zebrafish model for inflammation, in combination with scRNA-sequencing, to explore the integrated and cell-specific mechanisms of GR action *in vivo*. Using transgenic zebrafish lines, we first confirmed that glucocorticoid treatment reduced migration of both macrophages and neutrophils towards a site of inflammation. Initial bulk RNA sequencing analysis revealed that glucocorticoids modulated the expression of nearly all inflammation-induced genes, highlighting the effective and ubiquitous nature of glucocorticoid treatment. The scRNA-seq analysis revealed that in several key cell types, namely macrophages, neutrophils and epidermal cells, there was a clear differentiation in the transcriptome in response to amputation. In both amputation-specific epidermal cells and macrophages sensitivity to glucocorticoids was impaired, while neutrophils retained their responsiveness. To further understand the mechanisms underscoring these distinct glucocorticoid responses, we correlate the expression of specific transcriptional co-regulators with the observed cell-specific changes in responsivity. Taken together, we present a comprehensive view of cell type-specific transcriptional programs involved in acute inflammation and glucocorticoid responsivity *in vivo*.

Previous work from our group and others clearly established that treatment with glucocorticoids reduces (by ∼50%) the migration of neutrophils towards the site of inflammation in the zebrafish tail fin amputation model, which we confirmed in the present study(Chatzopoulou et al., 2016; Faught and Schaaf, 2023; Mathew et al., 2007; Xie et al., 2019). This suppression is dependent on GR activation, which reduces the amputation-induced *cxcl18b* and *il8* expression, which are known chemokine attractants of neutrophils in zebrafish(Xie et al., 2019). In the present study, glucocorticoid treatment also resulted in a small reduction in the migration of macrophages. In previous studies, the effect of glucocorticoids on macrophage migration was not as clear as the effect on neutrophils, likely because this migration is at least partially dependent on activation of MR(Faught and Schaaf, 2023; Xie et al., 2019). Here, we used the glucocorticoid dexamethasone to activate GR, and its ability to reduce macrophage migration may reflect the ability of this compound to also activate MR(Koning et al., 2022). Indeed, the more specific GR agonist beclomethasone did not affect macrophage migration (likely due to an inability to reduce the expression of genes encoding macrophage-specific chemo-attractants (i.e., *ccl2* and *cxcl11aa*)). However, beclomethasone did affect the transcriptional response and phenotypic signature of the macrophages, thereby inhibiting the differentiation of the macrophages towards a pro-inflammatory M1 phenotype(Xie et al., 2019). The responsivity at the level of the neutrophils in response to glucocorticoids has not yet been examined in this model.

It has been well described, in the zebrafish tail fin amputation model as well as in other models, that glucocorticoids, upon an inflammatory stimulus, reduce the expression of pro-inflammatory genes, and upregulate the expression of anti-inflammatory genes(Chatzopoulou et al., 2016; Xie et al., 2019). However, our bulk RNA-seq dataset revealed a far more nuanced scheme of how glucocorticoids modulate the inflammatory response. In line with our previous observations(Chatzopoulou et al., 2016; Xie et al., 2019), genes that showed strong regulation by amputation displayed a clear inhibition of this regulation by glucocorticoids. However, gene clusters with milder regulation by amputation showed an additive effect of amputation and glucocorticoid treatment. Notably, these clusters contained both pro-inflammatory and anti-inflammatory genes, suggesting that glucocorticoids do not universally suppress immune function. In human macrophages and microglia and glucocorticoids sensitize innate immune cells to specific stimuli by increasing NLR family pyrin domain containing 3 (NLRP3) inflammasome activity allowing for enhanced release of pro-inflammatory cytokines(Busillo et al., 2011; Busillo and Cidlowski, 2013; Feng et al., 2019). In line with these mechanistic studies, we also found upregulation of key inflammasome components, such as the NOD-like receptor (NLR) signaling pathway. Both the functional relevance and the mechanisms by which glucocorticoids induce the expression of pro-inflammatory genes are not yet well understood. It is thought that the apparently opposite effects appear to work together as a defense mechanism to both prepare the immune system and suppress it to restore homeostasis(Cruz-Topete and Cidlowski, 2014). Importantly, we did not observe any substantial gene cluster that was solely affected by amputation but not by DEX. These results are consistent with earlier data in zebrafish macrophages where almost all amputation-induced genes were affected by DEX(Xie et al., 2019). Taken together, DEX had a pervasive role in our dataset, enhancing weak but suppressing strong amputation-induced transcriptional effects.

The results from the bulk RNA-seq analysis did not provide any information on cell type-specific effects, but in examining our scRNA-sequencing data set it was immediately evident that there were amputation-specific cell clusters of epidermal cells, macrophage and neutrophils. The amputation specific epidermal cell population was highly enriched for many of the inflammation-induced genes. Based on its marker gene profile *(fgf20a*, *sp9*, *lef1*, and *il11a*(Chen et al., 2025; Hou et al., 2020; Poss et al., 2000; Whitehead et al., 2005)), pathway enrichment, and the relatively low cellular abundance, we postulate that this population corresponds to the wound epidermal cells localized at the distal side of the wound(Poleo et al., 2001; Poss et al., 2003; Tal et al., 2010; Gault et al., 2014; Pfefferli and Jaźwińska, 2015). Following tail fin amputation, a wound epithelium forms within approximately 30 min through collective migration of epidermal cells. Over the subsequent several hours, this tissue differentiates into the specialized apical epithelial cap, which is generally established by approximately 6 hpa and provides essential signals for blastema formation and regeneration(Pfefferli and Jaźwińska, 2015; Sehring and Weidinger, 2022). The wound epidermal gene expression at 4 hpa was furthermore characterized by high levels of inflammatory signaling molecules, tissue remodeling enzymes, and key metabolic hormones, including glucagon a (*gcga*) and leptin b (*lepb*). These latter genes suggest a metabolic shift to glycolysis and lipolysis respectively, and show that metabolic regulation is a central component of the acute inflammatory response(Sinclair et al., 2021),(Kang et al., 2016; Wu et al., 2024). Together, these transcriptional changes are may be aimed at controlling inflammatory processes, providing metabolic and structural resources for blastema formation, and facilitating extracellular matrix remodeling to enable tissue dedifferentiation and regeneration.

Importantly, glucocorticoid treatment did not abolish the formation of the wound epidermis in our study, indicating that formation of the wound epidermis *per se* is not targeted by glucocorticoids. However, we did observe glucocorticoid-induced changes in gene expression in the cells of the wound epidermis, such as downregulation of several regeneration-associated genes, encoding e.g., matrix remodeling enzymes, follistatin-like 3, and interleukin-11. The repression of these genes suggests that glucocorticoids, while allowing the formation of the wound epidermis, impair the function of this structure as a signaling hub to stimulate tissue regeneration.

Although the formation of the wound epidermis was not affected by DEX, the amputation-specific epidermal cells showed reduced glucocorticoid responsiveness. This is surprising, given the central role of the wound epidermis in chemotactic signaling. However, it should be noted that this reduced responsiveness was only observed with regard to GR-mediated transactivation. Indeed, our data suggests that increased expression of genes encoding specific components of the SWI/SNF complex (e.g., certain *smarc* paralogues), are negatively correlated with GR activity. The upregulation of these genes suggests enhanced ATP-dependent chromatin remodeling in these cells, indicative of dynamic reorganization of chromatin structure. This may suggest that the decreased responsivity of wound epidermal cells to GR activation is due to limited chromatin accessibility, a concept that has been thoroughly explored in the context of immune cells(Deochand et al., 2024; Oh et al., 2017). Taken together, our results suggest that inflammation alters the chromatin landscape in wound epidermal cells in such a way that suppression of gene expression by GR remains intact (e.g., of genes involved in chemotactic signaling(Xie et al., 2019), but that GR transactivation is reduced.

Besides the epidermal cells, we analyzed the leukocyte populations present in our dataset in terms of their transcriptional response to amputation and their glucocorticoid responsiveness. We identified, for both macrophages and neutrophils, a specific subpopulation that was only present upon amputation, suggesting that these subpopulations represent the neutrophils and macrophages that have responded to the amputation and migrated to the wounded area(Chatzopoulou et al., 2016; Xie et al., 2019; Ellett et al., 2011; Renshaw et al., 2006). Interestingly, the amputation-specific macrophages are characterized by a strong lipid metabolism signature, which is in line with an apparent increase in the activity of the Liver X Receptor alpha (LXRα, encoded by *nr1h3*), and enriched expression of its obligate dimerization partner, the Retinoic X Receptor Alpha (RXRα, encoded by *rxraa*). LXRα is a nuclear receptor activated by oxysterols, which are abundant in inflamed tissue(Brahmi et al., 2025; Carre et al., 2025), and are also generated within macrophages from lipids acquired during efferocytosis of apoptotic cells(Viaud et al., 2018). In macrophages, activation of LXRα induces a transcriptional program primarily aimed at cholesterol efflux and metabolic pathways to prevent lipid accumulation, but it is also exerts anti-inflammatory effects(Schulman, 2017; Leussink et al., 2020; Yan and Horng, 2020), driving macrophages from a pro-inflammatory (M1) to an anti-inflammatory, pro-resolving (M2) phenotype(Kim et al., 2016; Kimura et al., 2016). It has been shown that at 4 hours after the tail fin amputation in zebrafish larvae the macrophages at the wound site are in the transition from an M1 to an M2 phenotype(Xie et al., 2019; Nguyen-Chi et al., 2017; Begon-Pescia et al., 2026), and the data presented here suggest that this phenotypical shift is largely induced by oxysterol-induced activation of LXRα.

Similarly to the amputation-specific epidermal cells, the amputation-specific macrophages exhibited a reduced responsiveness to GR-induced gene activation compared to other macrophages. This may result from the observed differences in their coregulator profile, but may also be due to the increased LXRα activity in these cells. LXRα/RXRα heterodimers have been reported to suppress GR-mediated gene transactivation in mammalian cells *in vitro* and *in vivo*, by occupying glucocorticoid response elements and thereby preventing GR binding to these sites(Nader et al., 2012). A similar LXRα-mediated mechanism underlying reduced glucocorticoid responsiveness might occur in the amputation-specific macrophages in our study.

Interestingly, the pattern observed for neutrophils was markedly different. In recent years, increasing evidence has highlighted the considerable heterogeneity of neutrophils(Ng et al., 2019; Ganesh and Joshi, 2023; Riaz and Sohn, 2023). Consistent with this, we identified an amputation-specific neutrophil subpopulation displaying a pronounced pro-inflammatory phenotype. This population was enriched for genes involved in reactive oxygen species (ROS) production and chemotactic migration, and high expression levels of genes encoding pro-inflammatory transcription factors such as *rel* and *relb*. Thus, whereas macrophages appear to be transitioning towards an anti-inflammatory phenotype at this stage, neutrophils remain predominantly pro-inflammatory.

Furthermore, upon DEX treatment the neutrophil migration shows an approximately 50% reduction in migration towards the wound, while this treatment does not affect the size of this pro-inflammatory amp-specific neutrophil subpopulation. These findings suggest that differentiation into this pro-inflammatory state and migration towards the inflammatory site are not necessarily coupled processes, but are instead likely regulated by distinct signals originating from epidermal cells, macrophages, or both. In contrast to the amp-specific epidermal cells and macrophages, the amp-specific neutrophils showed no alteration in their glucocorticoid responsiveness, and showed transactivation and transrepression activity similar to the other neutrophils. Instead, they displayed glucocorticoid-induced transactivation and transrepression responses comparable to those of the remaining neutrophil populations. Although DEX treatment neither changed the size nor the overall identity of the amputation-specific neutrophil cluster, it did induce substantial transcriptional changes within these cells. This observation is consistent with the well-established glucocorticoid responsiveness of neutrophils(Hirsch et al., 2012; Ronchetti et al., 2018; Saffar et al., 2011) and may also have contributed to the reduced migration of this leukocyte population observed in the present study.

In summary, our analysis of injured and glucocorticoid-treated zebrafish caudal fin tissue revealed cell type-specific glucocorticoid responsiveness *in vivo*. Wound epidermal cells and LXRα-active macrophages exhibited reduced GR responsiveness, while activated neutrophils retained their responsivity. Reduced GR responsiveness was associated with an increased expression of SWI/SNF genes in wound epidermal cells, and increased activity of LXRα in amputation-specific macrophages. Development of novel GR ligands that target these coregulators and transcription factors, might provide a strategy to fine tune glucocorticoid responsiveness in a cell type-specific manner, potentially reducing adverse effects associated with systemic glucocorticoid therapy.

---

**Figure S1.**
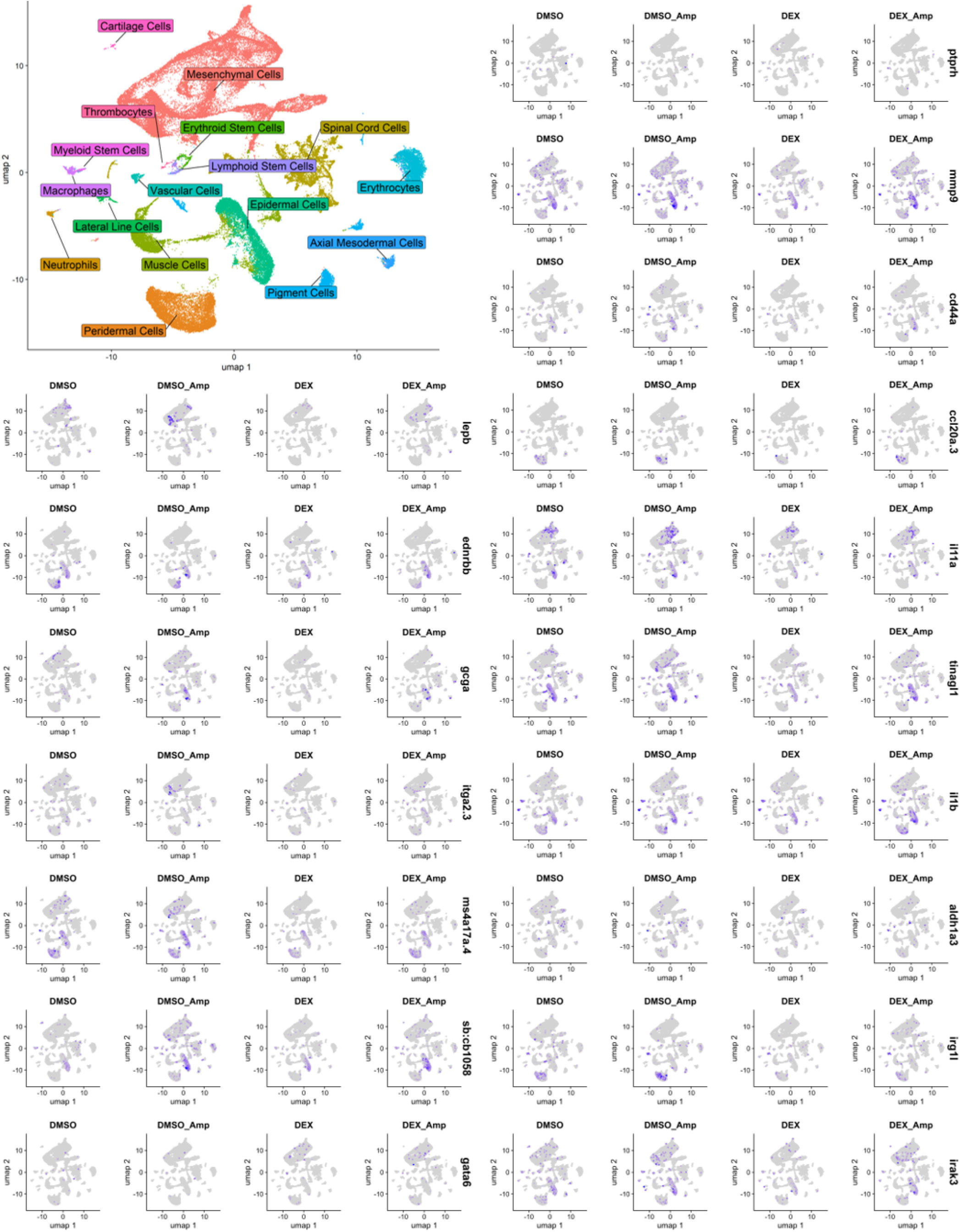
Feature plots of the top 17 genes found upregulated by amputation in bulk RNA-seq. Uncharacterized genes and genes that were not detected by scRNA-seq were excluded from this overview.

**Figure S2.**
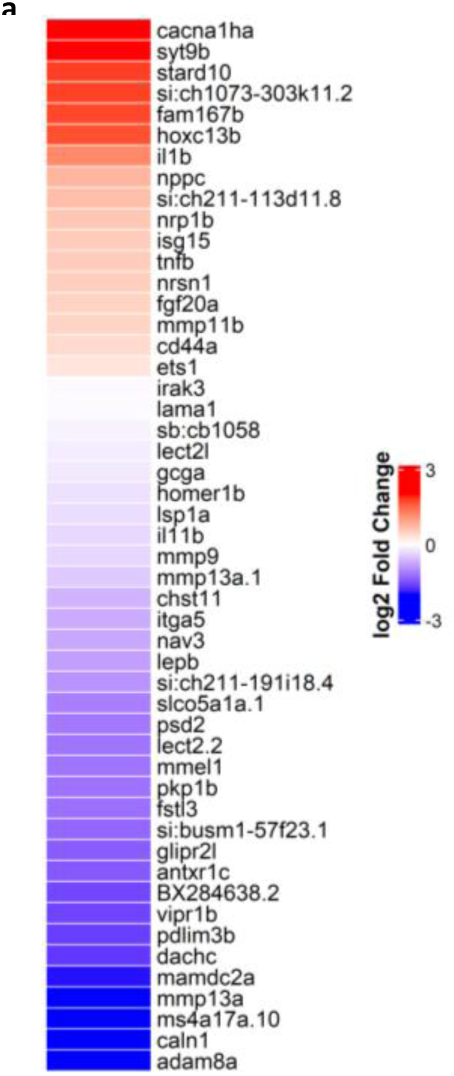
(a) Top 50 marker genes of the epidermal Amp-specific cluster, sorted by the DEX effect.

**Figure S3.**
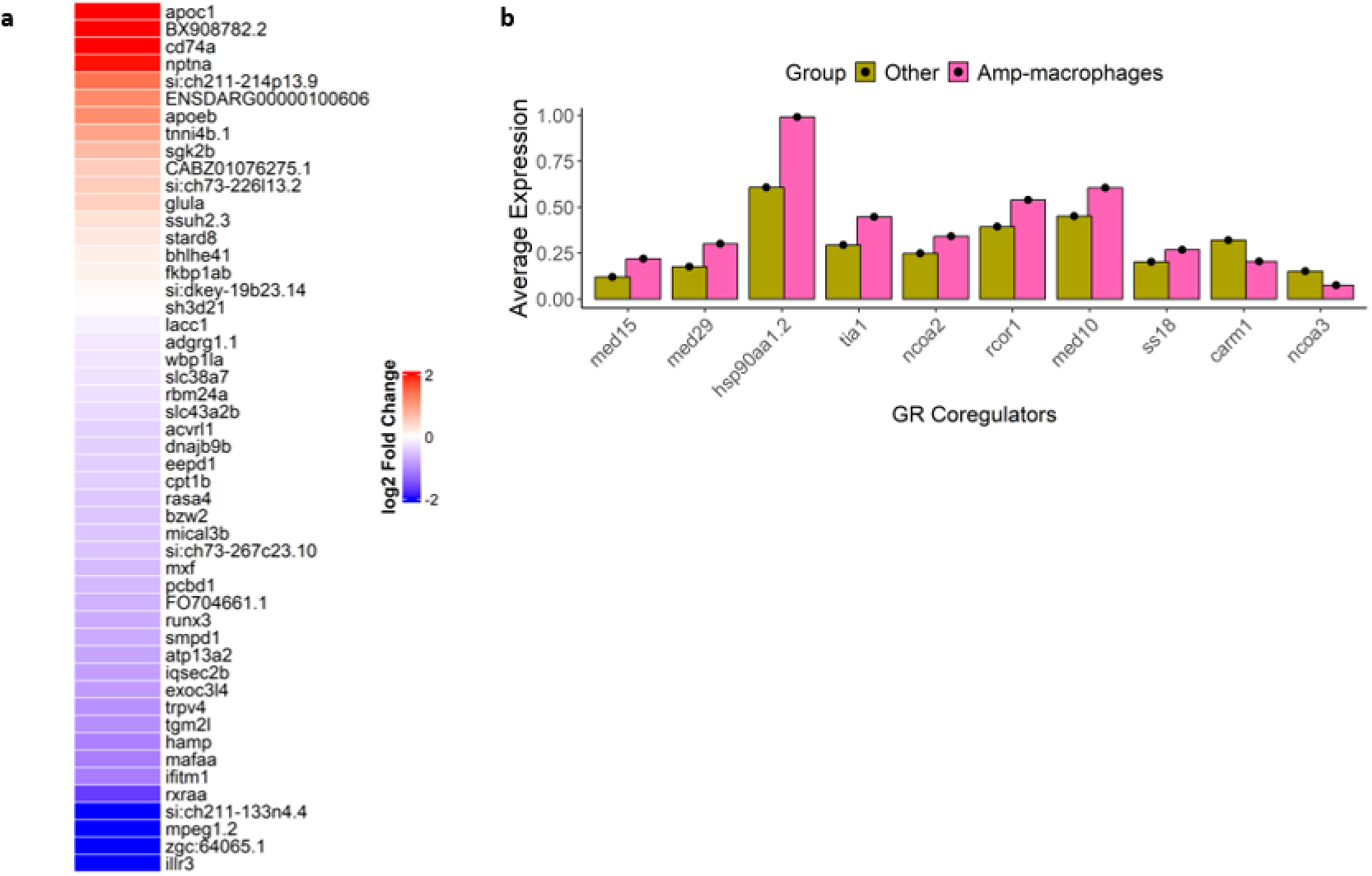
(a) Top 50 marker genes of the Amp-macrophages, sorted by the DEX effect. (b) Expression of GR coregulators in Amp-macrophages and other macrophages with a log2 fold change difference > 0.5.

**Figure S4.**
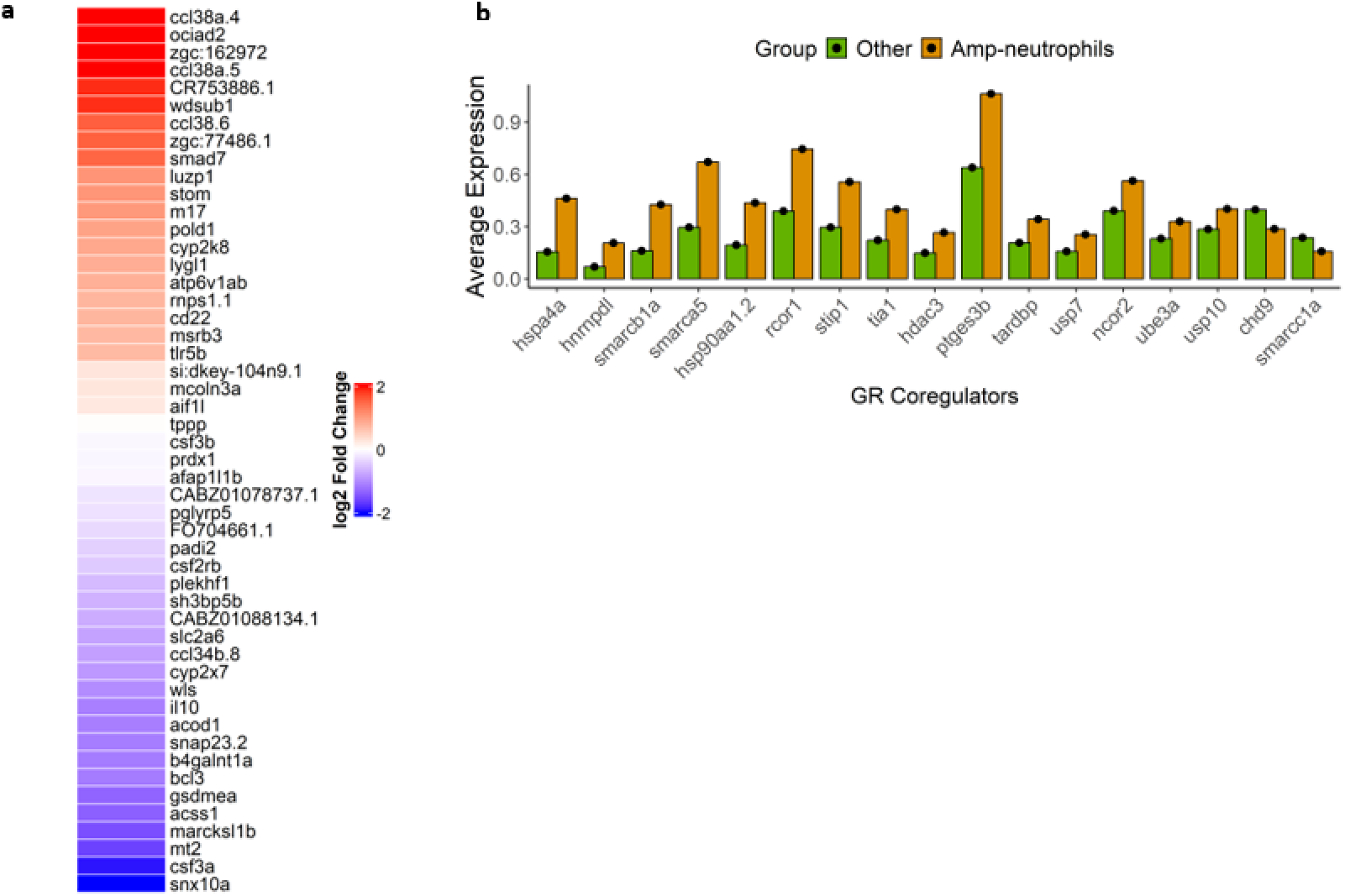
(a) Gene ontology analysis of Amp-neutrophil marker genes, sorted by the DEX-effect. (b) Expression of GR coregulators in Amp-neutrophils and other neutrophils with a log2 fold change difference > 0.5.

## METHODS

### Zebrafish lines and maintenance

Zebrafish (*Danio rerio*) were maintained and handled according to the guidelines from the Zebrafish Model Organism Database (http://zfin.org) and in compliance with the directives of the local animal welfare committee of Leiden University. Zebrafish were exposed to a 14 h light and 10 h dark cycle to maintain circadian rhythmicity. Fertilization was performed by natural spawning at the beginning of the light period. Eggs were collected and raised at 28°C in egg water (60 μg/ml Instant Ocean sea salts and 0.0025% methylene blue). In this study, the AB/TL wild-type zebrafish line was used.

### Caudal fin amputation in zebrafish larvae

In the caudal fin amputation experiments, 3 dpf larvae were pretreated with vehicle (0.05% DMSO) or 30 μM dexamethasone (DEX, Sigma-Aldrich, St. Louis, MO, USA) for 2 h before anesthetization in egg water containing 0.02% buffered aminobenzoic acid ethyl ester (tricaine; Sigma-Aldrich). Larvae were placed in Petri dishes coated with 2% agarose under a Leica M165C stereomicroscope (Leica Microsystems, Wetzlar, Germany), and the tails were amputated posterior to the tip of the notochord using a 1 mm sapphire blade (World Precision Instruments, Sarasota, FL, USA). The larvae were washed to remove the anesthetic and the treatment was continued for 4 h, after which the larvae were euthanized with an overdose of tricaine. Each experiment included non-wounded control groups treated for 6 h with DMSO or DEX.

### RNA sequencing analysis

For RNA sequencing (RNA-seq) analysis zebrafish larvae (120 larvae per condition) underwent the caudal fin wounding assay. After euthanasia with an overdose of tricaine, the caudal fin tissue halfway between the yolk extension and the fin tip was collected on a Petri dish coated with 2% agarose using a 1 mm sapphire blade. To block transcription, 1 μg/mL Actinomycin D (Focus Biomolecules, Plymouth Meeting, PA, USA) was added to the wash buffer. The tail tissue was collected on a 70 μm cell strainer and washed with PBS to remove debris. The tail tissue was subsequently transferred to a 15 mL centrifuge tube and spun down for 5 min at maximum speed. All liquid was removed, and the tissue was stored at -80°C until further use. The experiment was performed independently four times. Total RNA was extracted from the caudal fin tissue samples using TRIzol reagent (Thermo Fisher Scientific, Waltham, MA, USA) according to the manufacturer’s instructions. After DNase I treatment, a second TRIzol extraction was performed to remove DNase I and buffer salts. 10 μg RNA-grade glycogen (Thermo Scientific) was added during isopropanol precipitations to increase the recovery of low-concentration RNA from the solution. mRNA purification from total RNA, library preparation and sequencing were performed at GenomeScan (Leiden, Netherlands). Illumina NovaSeq 150 bp paired-end sequencing was performed with 20 million reads per sample. The resulting Fastq files were aligned to the zebrafish genome (GRCz11) with Rsubread 2.20, and features were counted using the *Danio rerio* annotation GTF 11.114(Liao et al., 2019). Differential gene expression values were determined by DESeq2 with standard settings(Love et al., 2014). Genes with an adjusted p-value lower than 0.05 according to the Benjamini-Hochberg procedure were considered significant. Heatmaps were prepared with the R package ComplexHeatmap(Gu et al., 2016). Pathway analysis was performed using clusterProfiler gseKEGG (ClusterProfiler::enrichKEGG, organism = “dre”, keyType = “ncbi-geneid”, pAdjustMethod = “BH”, pvalueCutoff = 0.1, minGSSize = 3, maxGSSize = 500, eps = 0)(Yu et al., 2012). Pathway analysis was run separately for six differentially regulated gene clusters and the combined results were visualized with ggplot2.

### Single-cell RNA sequencing sample preparation

For single-cell RNA sequencing (scRNA-seq) analysis, 3 dpf larvae (100 larvae per condition) underwent the caudal fin amputation assay, euthanasia and caudal fin tissue collection as before. The tail tissue was dissociated for 30 min at 30°C with 0.5 mg/mL Liberase TL (Roche, Basel, Switzerland) in PBS and trituration every 5 min with a P1000 micropipette. The cell suspensions were filtered once on a 70 μm and twice on a 30 μm cell strainer, and washed with PBS + 0.5% BSA. The filtrate was centrifuged for 5 min at 250 x g, and the cell pellet was resuspended in 100 μL PBS + 0.5% BSA. Sample viability above 85% was confirmed with Trypan blue using a TC20 cell counter (Bio-Rad). For each condition, approximately 30,000 cells were loaded on a Chromium X (10X Genomics, Pleasanton, CA, USA) and cell-barcoded libraries were prepared according to the manufacturer’s protocol for the Chromium GEM-X Single Cell3 kit v4 (CG000731 Rev B). Sequencing was performed on an Illumina NovaSeqX 10B 100 cycle flow cell (Illumina) reaching a depth of 25,000 reads per cell.

### Alignment of single-cell sequencing data

Alignment and processing of raw sequencing data were performed with 10x Genomics Cell Ranger v9.0.0 at the ALICE High Performance Computing facility of Leiden University. A primary *Danio rerio* genome assembly was prepared by the cellranger::mkref function from GRCz11 and the GTF 11.114. This reference was used to create a count matrix from the raw sequencing data using the cellranger::count functionality.

### Quality control and pre-processing

Filtered count matrices were loaded in Seurat v5 with parameters min.cells = 3 and min.features = 200(Hao et al., 2024). Each sample was treated separately but followed the exact same steps. We determined the mitochondrial gene content (Seurat::PercentageFeatureSet, pattern = “^mt-”) and ribosomal gene content (Seurat::PercentageFeatureSet, features = ribosomal genes). Cells with mitochondrial gene content > 10% were removed. Doublets identified by scDblFinder (artificialDoublets = 30000) were filtered out(Germain et al., 2022). Raw count data were log-normalized (Seurat::NormalizeData, normalization.method = LogNormalize, scale.factor = 10000, margin = 1). Cell cycle scoring was performed using Seurat v5 as based on *Danio rerio* cell cycle genes. Data from the four different conditions were merged (Seurat::merge) and layers were split. Data were scaled with regressing out mitochondrial gene content, ribosomal gene content, and the cell cycle (Seurat::ScaleData, vars.to.regress = c (“percent.mt”, “percent.ribo”, “S.Score”, “G2M.Score”)). Principal component (PC) analysis was performed (Seurat::RunPCA, npcs = 50), neighbors identified (Seurat::FindNeighbors, dims = 1:50, reduction = “pca”), biological similar cells were clustered together using the Leiden algorithm (Seurat::FindClusters, resolution = 1, algorithm = 4) and Uniform Manifold Approximation and Projection (UMAP) dimensionality reduction was performed (Seurat::RunUMAP, dims = 1:50, reduction = “PCA”). The data from the four conditions were integrated using Seurat’s RPCA method (Seurat::IntegrateLayers, method = RPCAIntegration, Seurat::JoinLayers, Seurat::ScaleData, and Seurat::RunPCA) so that biologically similar cells across the samples would cluster together irrespective of treatment. The optimal number of PCs was determined using a mathematical approach where the optimal number of PCs is equal to a 0.1% increase in variation compared to the previous PC.

### Annotation and interpretation of cell clusters

To annotate the identified clusters, clustering resolution was determined such that only the main clusters, assumed to be distinct tissues, would be separated. A combination of the clustree package and UMAP plots was used to determine the optimal resolution 0.3(Zappia and Oshlack, 2018). Seurat::FindAllMarkers was employed to identify marker genes per cluster with the parameters min.pct.1 = 0.1 and logfc.threshold = 2, unless otherwise indicated. Non-significant (p_val_adj < 0.05) markers were removed. The Daniocell database and the Zebrafish Information Network (ZFIN) were consulted to annotate the tissues based on the identified marker genes(Sur et al., 2023). After an initial low-resolution clustering, epidermal cell and leukocytes were subset for high-resolution clustering. For each subcluster, the most prominent marker genes were identified (Seurat::FindAllMarkers) and used for annotation. Cell annotations were transferred to the Seurat object and added as a new column in the metadata. UMAP reduction plots showing the cell clusters were created with Seurat::DimPlot, feature plots with Seurat::FeaturePlot, and dot plots with Seurat::DotPlot.

The Gene Ontology (GO) of marker genes (with log2 fold change > 2 for amputation-specific epidermal cells, and log2 fold change >1 for Amp-macrophages and Amp-neutrophils) was assessed with clusterProfiler (clusterProfiler::enrichGO, OrgDb = org.Dr.eg.db, ont = “BP” keyType = “ENTREZID”, pAdjustMethod = “BH”, pvalueCutoff = 0.05, qvalueCutoff = 0.05) with all expressed genes in the cell population as the universe background. The top GO terms by fold enrichment with gene count > 5 were subsequently visualized using ggplot2.

### Transcription factor activity inference

Transcription factor activity was inferred with decoupleR(Badia-i-Mompel et al., 2022). To date, no comprehensive transcription factor network database is available for zebrafish; therefore, a human database was used (OmnipathR:: get_dorothea (organism = “human”, levels = c (“A”, “B”, “C”))). To convert the human database to zebrafish, a gene_ortholog file from NCBI were used, and gene symbols were temporarily mapped to ENTREZ IDs via BiomaRt(Durinck et al., 2005; O’Leary et al., 2024). The zebrafish dataset was then used by decoupelR::run_mlm to infer transcription factor activity at single-cell level. The results were added to the main Seurat object (Seurat::CreateAssayObject) for downstream analysis (scaling and dimensional reduction). Differences in transcription factor activity between cell groups were assessed with Seurat::AverageExpression (assay = “TF_activity”) and visualized with ggplot2.

### GR Module Score determination, GR coregulator correlation and expression

For the GR responsiveness analysis, a GR Module Score (GMS) was assigned to every epidermal cell, neutrophil and macrophage. First, up- or downregulated genes were calculated per cell group (epidermis, neutrophils, and macrophages) with Seurat:FindMarkers, ident.1 = c (“DEX_Amp”, “DEX”), ident.2 = c (“DMSO_Amp”, “DMSO”), logfc.threshold = 0.5, min.pct = 0.1, test.use = “wilcox”. Next, the top 100 up- and downregulated genes by adjusted p-value were used to determine the average GR activity per cell using the Seurat::AddModuleScore function(Tirosh et al., 2016). Statistical differences in the GMS between different cell groups were assessed using a Wilcoxon test, and the results were visualized with ggplot2. To gain insight into genes that contribute most to the GMS difference between cell groups, the difference in average expression of all 100 genes between these cell groups was calculated and visualized. The GMS was subsequently correlated with expression of GR transcriptional coregulators. First, a list of coregulators was retrieved from the literature (Clarisse et al., 2017; Fadel et al., 2023), and filtered for presence in our dataset. Second, Spearman correlations between the GMS and expression of each coregulator were calculated for each cell. Only correlations with |ρ| > 0.2 for at least one of the two cell types (amputation-specific and other) were plotted. Statistical differences between the mean correlation values of the two cell types were determined by a Fisher Z-transformation and Benjamini-Hochberg procedure. Differences in coregulator expression between the cell types of interest were determined with Seurat:FindMarkers.

## ACKNOWLEDGEMENTS

We would like to acknowledge the animal caretakers at Leiden University for maintenance of our fish lines.

## COMPETING INTERESTS

No competing interests declared

## FUNDING

This work was supported by an NWO Veni grant (E.F) and the Biotech Boster Program (M.J.M.S).

## DATA AND RESOURCE AVAILABILITY

All datasets will be made available upon reasonable request

## AUTHOR CONTRIBUTIONS STATEMENT

Conceptualization (H.B., M.J.M.S., and E.F.); Methodology (H.B.); Investigation and Formal Analysis (H.B., T.S., A.W., N.E., and S.K.); Data curation (H.B., M.J.M.S., and E.F.); Writing-original draft (H.B.), Writing – review and editing (H.B., M.J.M.S., E.F., and A.M.).

## Notes

### Competing Interest Statement

The authors have declared no competing interest.

